# A global test of hybrid ancestry from genome-scale data

**DOI:** 10.1101/2023.02.24.529943

**Authors:** Md Rejuan Haque, Laura Kubatko

**Author notes:** **Corresponding author: Md Rejuan Haque,**. **Laura Kubatko,**.

## Abstract

Methods based on the multi-species coalescence have been widely used in phylogenetic tree estimation using genome-scale DNA sequence data to understand the underlying evolutionary relationship between the sampled species. Evolutionary processes such as hybridization, which creates new species through interbreeding between two different species, necessitate inferring a species network instead of a species tree. A species tree is strictly bifurcating and thus fails to incorporate hybridization events which require an internal node of degree three. Hence, it is crucial to decide whether a tree or network analysis should be performed given a DNA sequence data set, a decision that is based on the presence of hybrid species in the sampled species. Although many methods have been proposed for hybridization detection, it is rare to find a technique that does so globally while considering a data generation mechanism that allows both hybridization and incomplete lineage sorting. In this paper, we consider hybridization and coalescence in a unified framework and propose a new test that can detect whether there are any hybrid species in a given set of species. We propose that based on this global test of hybridization, one can decide whether a tree or network analysis is appropriate for a given data set.

## 1 Introduction

Advances in modern sequencing technologies have led to the availability of a substantial amount of DNA sequence data from hundreds, if not thousands, of species. These data are often used to understand the evolutionary relationships between species as well as the process of evolution more generally. One way of representing the evolutionary relationships between a collection of species is with a species-level phylogeny, also known as a phylogenetic tree. A species-level phylogeny is an acyclic graph in which all edges are considered to be directed from the root node to the leaves. An internal node, at which a population splits into two, with each descendant evolving independently, represents a speciation event. In a phylogeny, all internal nodes except the root node are connected to three edges, making the degree of all internal nodes three. The root node, which represents the common ancestor of all the species in the data, is connected with two edges leading the root node to have degree two, while the leaves (also known as taxa) have degree one and represent the present-day populations from which the DNA sequence data were collected (see right panel of Figure 1).

**Fig. 1:**
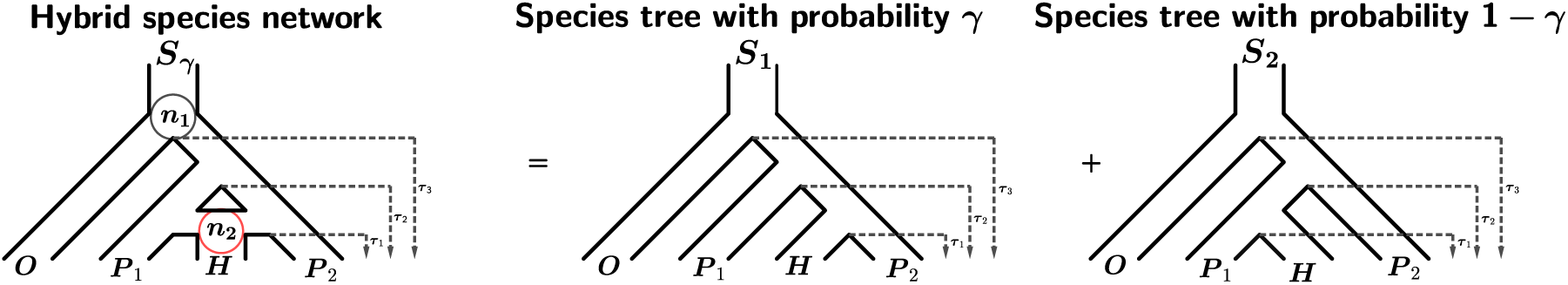
Model for the species-level relationships among four taxa under the coalescent model with hybridization. Here taxon *H* is a hybrid of taxa *P*_1_ and *P*_2_, *τ*_1_, *τ*_2_, and *τ*_3_ are the speciation times for the internal nodes from the present. The circles on the leftmost figure represent internal nodes. The red circle (*n*_2_) represents the hybrid node as there are three descendent from that node. The network on the left can be decomposed into two species-level phylogenies; one with *H* and *P*_2_ as sister taxa with probability *γ*, and one with *H* and *P*_1_ as sister taxa with probability 1 – *γ*.

One major drawback of using a phylogenetic tree to represent evolutionary history is that it assumes speciation occurs through bifurcation, meaning that an ancestor can have precisely two descendants. However, this might not always be the case since there are other evolutionary mechanisms that generate novel species. One such mechanism is hybridization, which occurs when two distinct populations interbreed and produce a new species that share genetic material with both parents. However, hybridization does not always result in creating a new species, in which case the mechanism is known as introgression [1–9]. Recent studies have shown numerous instances of hybridization in plants and animals, despite the belief that hybridization is rare [9–11]. If a collection of species are known to include species that have arisen via hybridization, one should infer a species network rather than a phylogenetic tree. Since the internal nodes in a network are allowed to have degree of more than three, networks can capture evolutionary processes such as hybridization (see left panel of Figure 1). However, inferring a network from data can be computationally intensive, especially when the number of taxa and/or the length of the DNA sequence alignment are large [12]. In addition, network analysis in the absence of hybridization can lead to incorrect conclusions concerning the ancestry of the taxa identified to of hybrid origin. Thus, an important first step before deciding whether to perform a network or a tree analysis is to gather concrete evidence about the possibility of hybridization among the species on which inference is to be made.

In the literature, methods for detecting hybridization are ubiquitous. However, several of these methods ignore a crucial evolutionary process called incomplete lineage sorting (ILS). In practice, the DNA sequence data come from genes, and each of these genes has its own evolutionary history, which may or may not match the evolutionary history of the species. When two gene copies from the current generation fail to coalesce in the immediately previous generation, incomplete lineage sorting (ILS) results, leading to gene-tree species-tree discordance [13, 38]. Coalescent theory links the gene trees to a species tree by providing the probability distribution of all possible rooted gene trees arising within a rooted species tree and thus can be used to model ILS [14–16]. To distinguish between hybridization and incomplete lineage sorting, Joly et al. (2009) proposed a statistic based on the observation that the genetic distance between two DNA sequences should be smaller in the presence of hybridization than the distance that would be observed if only ILS is acting on the genes. One can simulate this distance under the null hypothesis that ILS alone explains the discordance using coalescent theory. Once the distribution of the distance under the null hypothesis is estimated, one can compare the observed distance to the null distribution to assess the evidence for hybridization [17].

Tests based on the specific site pattern frequencies (a particular position at an alignment is called a site, and the combination of possible nucleotides at a particular site for the sampled individuals is called a site pattern) have also been developed to test ancient admixture utilizing Patterson’s D-Statistic [18–20]. In addition, several likelihood-based methods have been proposed, including Meng and Kubatko (2009), Kubatko (2009), and Yu et al. (2014). In 2019, Kubatko and Chifman proposed a method to detect hybridization based on “phylogenetic invariants” (polynomials in the site pattern probabilities that evaluate to zero on any probability distribution that is consistent with the tree topology and associated model). Their method considers models that can generate data that allows the possibility of hybridization as well as incomplete lineage sorting. Based on a comparison of popular hybridization detection methods, Kong and Kubatko (2021) concluded that Kubatko and Chifman’s (2019) invariant-based method detects hybrid species quickly and accurately, generally outperforming the other methods tested.

One common characteristic of several of these available methods is that they test whether a suspected species is a hybrid given knowledge of its two parents. However, to decide whether or not a network analysis is necessary, we need to perform a global test to assess evidence for the presence of one or more hybrid taxa in the whole data set. Kubatko and Chifman (2019) extended their method to deal with phylogenies containing more than four taxa by testing every species as a hybrid and applying a Bonferroni correction to adjust for multiple testing. However, the number of tests one needs to consider can be massive, especially when the number of taxa is large, making the Bonferroni correction less powerful in detecting hybridization events. For instance, from a data set containing *m* species, one can pick three species and test three hypotheses considering one of the three species as a hybrid and the other two as parents. Hence, one can test a total of 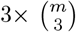 hypothesis using a data set containing *m* species. If, for instance, a data set contains eight species, we need to perform 168 tests, and the number of required tests grows significantly as the number of taxa increases. In addition, the individual tests are correlated because the test statistic for each individual test is calculated using sequences from closely related species, making the Bonferroni method conservative. Thus, to perform a global test for detecting hybridization, we need a method that can handle arbitrary correlation structures. One possibility is to use a combination test.

A combination test combines the individual p-values to aggregate multiple small effects rather than adjusting individual p-values to maintain the false rejection rate. Fisher’s combination test is the oldest method of combining individual p-values to test an overall null hypothesis [26]. Consider a scenario where we carry out *l* individual tests intending to determine whether all of the null hypotheses are true. Fisher’s combination test statistic to perform this test is as follows:

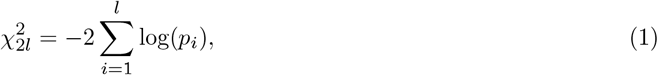

where *p_i_; i* = 1, 2,…, *l*, is the p-value from the *i^th^* individual test. The test statistic above converges to a Chi-squared distribution with *2l* degrees of freedom under the assumption that all individual nulls are true and the *p_i_* are independent. When the individual p-values are small, the value of the test statistic in Equation 1 tends to be larger, leading to evidence suggesting that not all of the null hypotheses are true. The test has substantial power to reject the global null hypothesis when a large fraction of the individual p-values are small. This will not be the case, however, when only a small fraction of the individual tests are expected to be significant [27, 28], a situation that we might often encounter in many applications of bioinformatics. For instance, while assessing genome-scale data to detect hybridization events for 20 taxa, one needs to perform a total of 2280 tests, only a few of which are expected to be significant. The application of Fisher’s combination test, in this case, may lead to failure to detect any hybrid species.

Multiple attempts have been made to improve the power of the test in situations like this. The most popular combination tests that attempt to improve power are Tippett’s minimum p-value [29] and higher criticism tests [30]. In Tippett’s minimum p-value approach, the test statistic is determined by the minimum of the individual p-values. That is, for the *l* individual p-values (*p*_1_, *p*_2_,…, *p_l_*), the test statistic is defined as: *S_T_* = *min*(*p*_1_, *p*_2_,…, *p_l_*). Assuming all the individual p-values are independent and all the individual nulls are true, Tippett’s minimum p-value statistic (*S_T_*) converges to a *Beta*(1, *l*) distribution. For *l* individual tests with *α* = 0.05, the higher criticism test statistic proposed by Tukey is

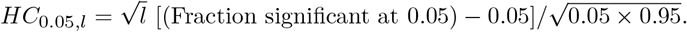

In Tukey’s interpretation, the null hypothesis should be rejected when *HC* is greater than 2, assuming that the individual p-values are independent. When the assumption of independence of individual p-values holds, Tippett’s minimum p-value and the higher criticism tests both have high power. However, both these tests, as well as Fisher’s combination test, fail to perform well when there is dependence among the individual tests.

In reality, it is often the case that the individual tests have a dependence structure and the p-values are correlated. For example, in a phylogenetic tree, species are related by their common ancestor. There are methods available based on permutation or numerical simulation to incorporate the correlation structure [31]. However, these approaches are computationally involved and time-consuming when performing a huge number of individual tests involving massive data sets. Recently, an analytical approximation method has been proposed using the higher criticism test to incorporate the dependence structure [32]. However, this approximation is computationally involved and does not perform well when the p-values are extremely small.

To incorporate the arbitrary dependence structure of individual tests and efficiently handle a large number of tests, Liu and Xie (2020) proposed a combination test called the Cauchy combination test. Following Fisher’s combination test, the test statistic of the Cauchy combination test is calculated using the weighted sum of transformed p-values. Let *p_i_* be the individual p-value, for *i* = 1, 2,…, *l*. Then, the Cauchy combination test statistic is defined as

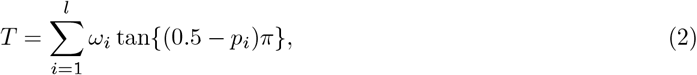

where the weights *ω_i_*’s are nonnegative and 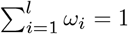. The authors showed that the statistic in Equation 2 converges to a standard Cauchy distribution under the global null hypothesis even when there is an arbitrary correlation structure between the individual p-values. The only assumption made by the authors is that any two of the individual test statistics should have a bivariate normal distribution. That is, considering *X_m_* to be the *m^th^* test statistic, *m* = 1, 2,…, *l*, for any 1 ≤ *i* < *j* ≤ *l*, (*X_i_, X_j_*)*^T^* follows a bivariate normal distribution. However, Liu and Xie (2020) argued that the bivariate normality assumption is mild, and the test is robust even when the assumption is not satisfied.

In 2022, Zhongxue Chen showed situations where the Cauchy combination test failed to reject the global null hypothesis even if some of the individual tests were rejected correctly. The author proposed a robust test combining the MinP [29] and the Cauchy combination [33] test that performs well under a wide range of situations. Consider combining *l* individual tests with p-values: *p*_1_, *p*_2_,…, *p_l_*. Let the p-value for the global null hypothesis using the Cauchy combination test be *p_c_*, and the p-value using the MinP test be *p_m_* with *p_m_* = *I* × *min*(*p*_1_, *p*_2_,…, *p_l_*). Using the resulting p-values from the Cauchy combination and MinP test, Chen (2022) proposed a test statistic called MinP-CCT-MinP (MCM) with which the p-value for the global test is calculated as follows:

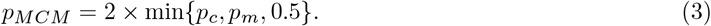

Chen (2022) showed that the proposed test is robust and performs well in situations where the Cauchy combination test might fail while maintaining the correct type I error rate. Finally, the author proved that unlike the Cauchy combination test, the MCM test does not need any distributional assumption.

Here, we consider the test statistic proposed by Kubatko and Chifman (2019) to test whether a particular species is a hybrid given two parents in both the Cauchy combination and MCM test framework proposed by Liu and Xie (2020) and Chen (2022), respectively, to suggest a global test statistic for detecting the presence of hybridization within a set of species. We will perform an extensive simulation study to investigate the performance of the proposed test. Finally, we will apply our method to empirical data sets and compare results with existing studies of those data sets.

## 2 Methods

As discussed in the previous section, ILS is a crucial evolutionary mechanism that can lead to gene-tree species tree discordance and thus should be incorporated into phylogenetic applications. Hence, we will consider a data generation mechanism that allows both hybridization and ILS, following the model proposed by Meng and Kubatko (2009). As already discussed, a bifurcating species-level phylogeny is inadequate to represent a hybridization event. Instead, a species network that connects two edges of a phylogenetic tree using a horizontal line is commonly used to depict a hybrid node. Consider the phylogenetic network in the leftmost panel of Figure 1. Here *H* is a hybrid species that contains a portion (*γ*) of its genetic material from *P2* and the rest (1 – *γ*) from parent *P*_2_; *τ_i_* is the speciation time for node *i* measured from the present.

We assume that the aligned DNA sequences arise along the network. Let *Y_L_* be the observed nucleotide for a specific site in the DNA sequence for taxon *L*. Then a site pattern arising from the species network in Figure 1 can be expressed as *Y_O_ Y*_*P*_1__ *Y_H_ Y*_*P*_2__. For a four-taxon network, the number of possible site patterns is 4^4^ = 256 since *Y_L_* ∈ {*A, G, C, T*}. Let *p_ijkl_* = *P_τ_*(*Y_O_* = *i*, *Y*_*P*_1__ = *j*, *Y_H_* = *k*, *Y*_*P*_2__ = *l*|*S_γ_, τ*)) be the probability of site pattern *ijkl* with *i, j, k, l* ∈ {*A, C, G, T*}, and let ***P*** = (*P_AAAA_*|(*S_γ_, τ*),…, *P_TTTT_*|(*S_γ_, τ*)) be the set of all possible site pattern probabilities on network (*S_γ_, τ*). If we let *N_ijkl_* be the observed number of sites with site pattern *ijkl* and *N* be the total number of sites in the aligned DNA sequence data, one can estimate *Pijki* as 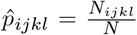, and the set of observed site pattern probabilities can be written as 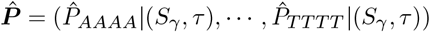.

As the vector 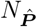 represents the observed count of each of the 256 possible site patterns, it is easy to see that

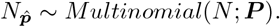

With *i* ≠ *j* ∈ {*A, G, C, T*} consider the following linear relationships:

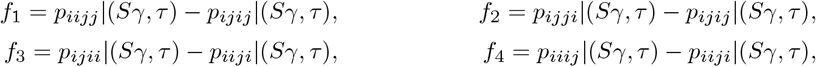

Kubatko and Chifman (2019) showed that *f*_2_ and *f*_4_ evaluate to zero assuming the site patterns arise from tree *S*_1_ (middle panel of Figure 1), while *f*_1_ and *f*_3_ are non-zero, and the opposite is true considering *f*_1_ and *f*_3_ with species tree *S*_2_ (rightmost panel of Figure 1). The authors also showed that 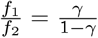, and using this relationship, they defined a test statistic called the Hils statistic to test *H*_0_: *γ* = 0:

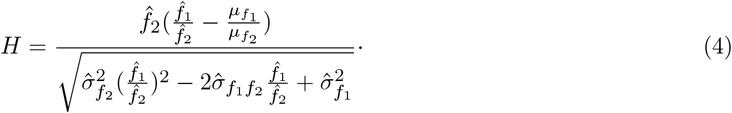

Here, *μ*_*f*_1__ and *μ*_*f*_1__ are the mean of *f*_1_ and *f*_2_, respectively, *σ*_*f*_1__ and *σ*_*f*_2__ are the standard deviation of *f*_1_ and *f*_2_, respectively, and *σ*_*f*_1_*f*_2__ is the covariance between *f*_1_ and *f*_2_. It has been shown that under *H*_0_, the statistic *H* follows a standard normal distribution and the null is rejected for large values of the observed statistic (Kubatko and Chifman, 2019).

Now, consider testing the following hypothesis given aligned DNA sequence data for *m*+1 taxa (*m* ingroup species and one outgroup):

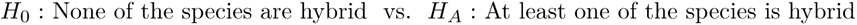

Consider choosing three ingroup taxa and the outgroup making a four-taxon subset from the set of taxa. Using the selected subset, we can consider a quartet network assuming that one of three ingroup taxa is a hybrid and the other two are the parental species. Note that we can build three such networks from one four-taxon subset by designating each ingroup taxon as the hypothesized hybrid species. For instance, if the selected four-taxon subset contains outgroup *O* and ingroup taxa *P*_1_, *P*_2_, and *H*, we can build three networks assuming *H* as a hybrid of parents *P*_1_ and *P*_2_, *P*_1_ as a hybrid of parents *H* and *P*_2_, and *P*_2_ as a hybrid of parents *H* and *P*_1_. Using the whole collection of species, one can build 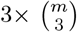 quartet networks with the outgroup and three taxa (triplet) selected from the *m* ingroup taxa. For instance, one can build a total of 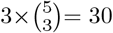 quartet networks from a collection of species containing one outgroup and five taxa.

Note that for a network with *m* taxa, *4^m^* site patterns are possible. To get the site patterns for a quartet network built by selecting a four-taxon subset from the collection of taxa, one can marginalize over the missing species. For instance, consider creating quartet networks from the species network (*S_γ_, τ*) in Figure 2a containing six species. Let the *k^th^* quartet network (*S_γ_k__*) contains species *O* as outgroup, *h* as hybrid, and *p*_1_, *p*_2_ as the parental species. Then the site pattern probability 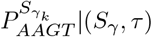 can be found by:

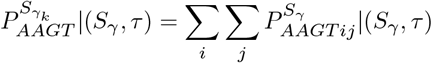

here, *i, j* ∈ {*A, G, C, T*} represents the possible site patterns of species *α* and *b* from network (*S_γ_, τ*) in Figure 2a that were not selected in the *k^th^* quartet network (*S_γ_k__*), respectively. It is clear that after marginalization, the possible number of site patterns for a quartet network is 4^4^ = 256.

**Fig. 2:**
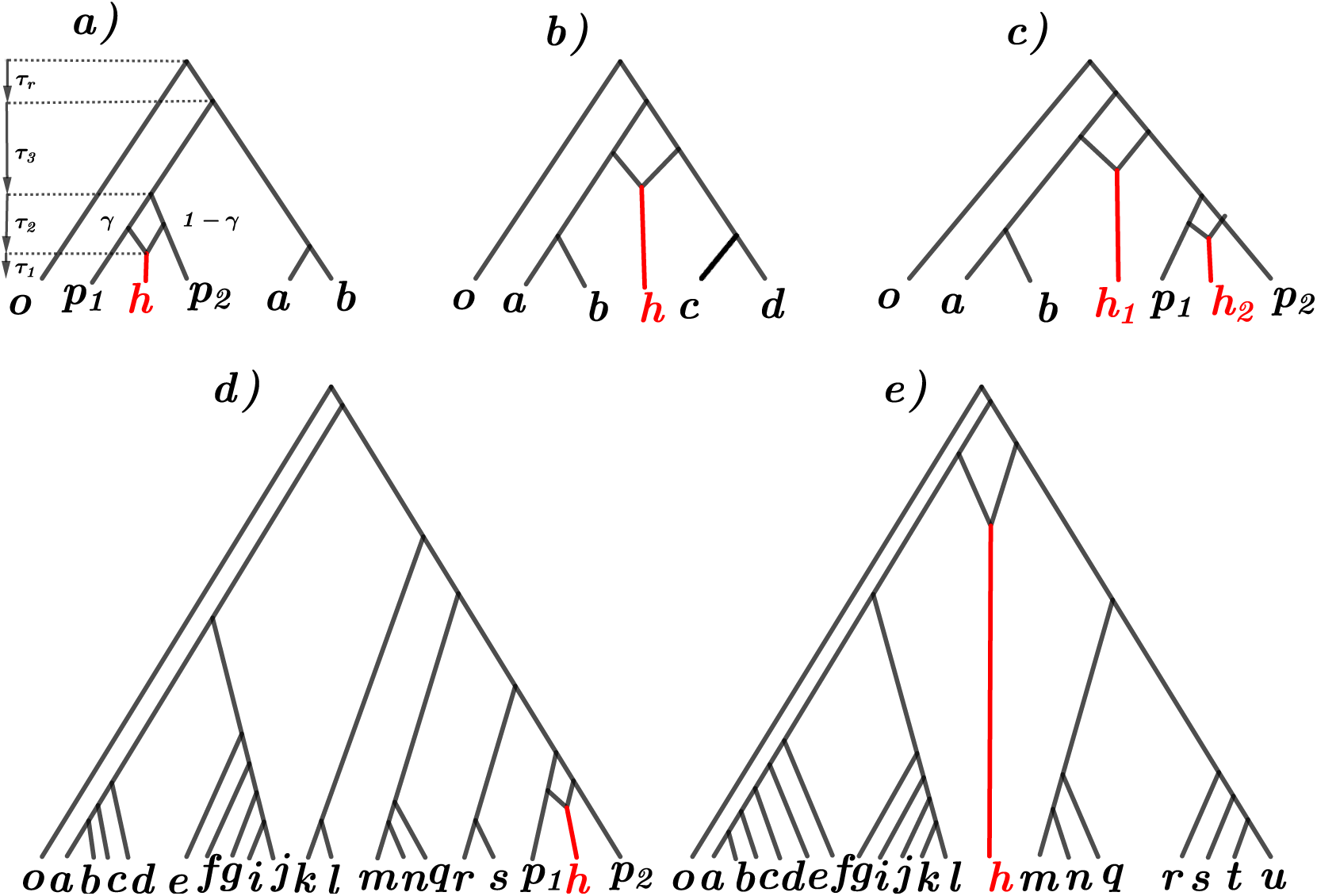
Five hybridization scenarios were considered in this study: a) one recent hybridization event within six species, b) one ancient hybridization event within six species, c) one recent and one ancient hybridization event within seven species, d) one recent hybridization event within twenty species, and e) one ancient hybridization event within twenty species. The outgroup species is denoted by *o*. For panels a, b, and d, the hybrid species is denoted by *h*, and parents of the hybrid species are denoted by *p*_1_ and *p*_2_. For panel c, the two hybrid species are denoted by *h*_1_ and *h*_2_; the parents of *h*_2_ are *P*_1_ and *p*_2_, and the parents of *h*_1_ are the ancestor of *a* and *b*, and the ancestor of *p*_1_, *h*_2_, and *p*_2_. For panel e, the hybrid species is denoted by h, and its parents are the ancestor of *a, b*,…, *l*, and the ancestor of *m, n*,…, *u*. *γ* is the proportion of genetic material shared by the leftmost parent to the hybrid. The time between speciation event *l* and *l* + 1 is denoted by *τ_i_* Note that only the first sub-figure (panel a) is fully labeled.

Once the set of four taxa for the *k^th^* 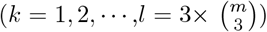 quartet network is selected, one can decompose the network (*S_γ_k__*) into two species trees (*S*_*k*1_ and *S*_*k*2_) following Figure 1. Also, following the idea of Kubatko and Chifman (2019), one can define 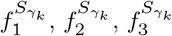, and 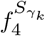 using the quartet network *S_γ_k__* and show that 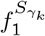 and 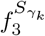 are zero when evaluated on the site pattern probabilities arising from one of the species trees (e.g., *S*_*k*1_) and not zero when evaluated on the site pattern probabilities arising from the other species tree (e.g., *S*_*k*2_). On the other hand, for 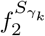 and 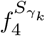, the opposite is true. However, none of the linear relationships are zero when evaluated on the site pattern probabilities from the hybrid species network *S_γ_k__* with *γ_k_* ∈ (0, 1). One can obtain a function of *γ_k_* using these linear relationships as follows:

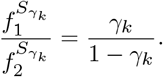

Now, one can easily estimate these functions by substituting the observed site pattern probabilities and then use the properties of the multinomial distribution and easily estimate the means 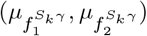 and variances 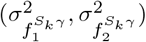 of these estimated functions. It is also possible to estimate the covariance 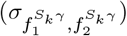 between these functions. Once the means, variances, and covariances are computed, one can use the *H* statistic in Equation 4 to test *H*_0_: = 0 for the *k^th^* quartet network (*S_γ_k__*):

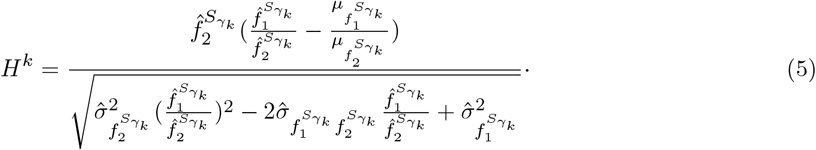

Under the null hypothesis that *γ_k_* = 0, the term 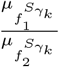 is 0. Also, note that under the null hypothesis, *H^k^* follows the standard normal distribution. Thus, the p-value of this test can be computed easily as follows:

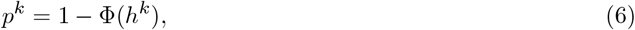

where *h^k^* is the observed value of the test statistic and Φ(*h^k^*) = *Pr*(*Z* < *h^k^*) with *Z* ~ normal(0, 1). Note that one can repeat this process and compute the p-values for all possible tests. For instance, for a species network with five ingroup and one outgroup species, one can compute a total of 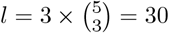 p-values. These tests will not be independent as species in a tree are related through their common ancestors. Let *p*^1^, *p*^2^,… *p^l^* be the corresponding p-values of the *l* tests. To test the global null hypothesis: *H*_0_: None of the species are hybrid, we propose the following statistic:

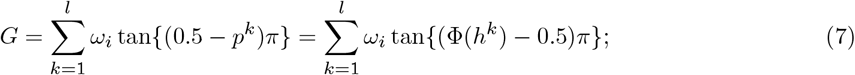

here, the *ω_i_*’s are non-negative weights such that 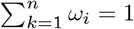 *ω_i_* = 1. Under the global null hypothesis *H*_0_, the test statistic *G* follows a standard Cauchy distribution [33]. Note that a large value of the test statistic *G* provides evidence against the global null hypothesis. Thus the c.d.f. of the standard Cauchy distribution can be used to compute the p-value for the global null as follows:

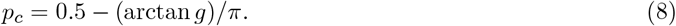

Here, *g* is the realized value of *G*. The global null hypothesis is rejected if the p-value is smaller than *α* given that the chosen significance level is *α*. Note that in applying the results of the Cauchy combination test [33], we have assumed that for any 1 ≤ *i* < *j* ≤ *l*, (*H^i^, H^j^*)*^T^* follows a bivariate normal distribution. The bivariate normality assumption is reasonable as the site pattern probabilities are well approximated by a normal distribution when the number of sites is large. Also, the summation of any two test statistics (e.g., *H*^1^ + *H^3^*) is just the sum of two approximate normal random variables, and thus the sum is also normal. Hence, it is safe to assume bivariate normality here.

Now, to apply the MCM proposed by Chen (2022), we will follow the following two steps:

- Step 1: Using the individual p-values (*p*_1_, *p*_2_,…, *p_l_*), calculate the p-value (*p_m_*) for the global test using the MinP test with *p_m_* = *l* × min{*p*_1_, *p*_2_,… *p_l_*}.
- Step 2: Calculate the p-value (*p_mcm_*) for the global test of no hybridization event using the MCM test with *P_mcm_* = 2 × min{*p_c_, p_m_*, 0.5}, where *p_c_* is the p-value for the global test from the Cauchy combination test.

### 2.1 Simulation Study

To investigate the performance of the proposed global test for detecting the presence of hybridization within a collection of species, we designed an extensive simulation study with five different species networks covering several evolutionary scenarios (Figure 2). We varied the number of taxa, the number of hybridization events, the time of hybridization, the branch lengths (*τ_i_*), the hybridization parameter (*γ*), the length of the simulated sequences, and the effective population size (*θ*) to generate DNA sequence data. To observe how the proposed method behaves when the data consist of a small number of taxa, we generated data using two networks with six taxa (Figure 2a and 2b).

In one of the six-taxon networks (Figure 2a), there is a recent hybridization event close to the leaf nodes, whereas the hybridization event occurs close to the root node in the other six-taxon network (Figure 2b) (often called ancient hybridization). Similarly, we have considered two twenty-taxon networks, one with recent hybridization (Figure 2d) and one with ancient hybridization (Figure 2e). Finally, we have considered a seven-taxon network with one recent and one ancient hybridization event (Figure 2c). For all five hybridization scenarios, the value of each *τ_i_* is set to be 0.5 or 1.0. The speciation time between the outgroup and the ingroup (*τ_r_*) is set at 5.0 to ensure that the genetic material between the two groups is sufficiently different. For each scenario, the effective population size *θ* has been fixed to 0.001, 0.005, 0.01, or 0.03. Eleven different values of *γ* are chosen (*γ* ∈ {0.00, 0.05, 0.10, 0.15, 0.20, 0.25, 0.30, 0.35, 0.40, 0.45, 0.50}) for each setup. Combining all these parameter values, we will have 88 scenarios in total for each of the networks presented in Figure 2.

We simulated a total of 1000 gene trees from each of the 88 different scenarios described above using *ms* [35]. Note that *ms* only accepts input in a tree format. Consequently, generating gene trees directly from a species network is thus impossible using *ms*. Species networks with hybridization, however, can be viewed as a combination of species trees since a network with *l* hybrid species can be decomposed into 2*^l^* species trees by making the hybrid a sister taxon to one of its parents (Figure 1). Once the desired network is decomposed into species trees, one can simulate gene trees from each of the species trees accordingly to achieve the desired *γ*. For instance, to achieve a *γ* = 0.5 from Figure 1, one can generate 500 gene trees using species tree *S*_1_ (middle panel of Figure 1) and 500 gene trees using species tree *S2* (rightmost panel of Figure 1). For a tree with two hybridization scenarios, one can use a total of four trees and can generate 250 gene trees from each of those four species trees to get 0.5 as the value of one of the *γ* (e.g., *γ*_1_) while fixing the other *γ* (e.g., *γ*_2_) at a fixed value 0.5. An example *ms* command to generate 500 gene trees and save it to a file named *tre.1* from the species tree *Si* (middle panel of Figure 1) is: *ms 4 500 -t 5 -T -I 4 1 1 1 1 -ej 0.5 4 3 -ej 1.0 3 2 -ej 5 1 2*

|*tail -n +4*| *grep -v > tre.1*.

To simulate the DNA sequence data from the network, we used the gene trees generated using *ms* as input to *seq-gen* [36]. The HKY model was used for DNA substitution with the ratio of transitions and transversions equal to 1(*κ* = 1), meaning our model assigns equal probability to transitions and transversions. Also, we set frequencies of nucleotide *A, C, G*, and *T* at 0.300414, 0.191363, 0.196748, and 0.311475, respectively, following the work of Yu et al (2014). The length of each of the sequences was set to be 50, 100, 150, and 200, which led to a total sequence length of 50 × 10^3^, 100 × 10^3^, 150 × 10^3^, and 200 × 10^3^, respectively, when the sequences were concatenated using *goalign* (https://github.com/fredericlemoine/goalign). Finally, the aligned DNA sequence data were converted to *Phylip* format, and a *map* file mapping individuals to the corresponding population was created. These two files were used to calculate the individual *H* statistic and the corresponding p-value.

The resulting data for each of the simulation setups, as well as a Python package to carry out the proposed test, can be found at https://github.com/rhaque62/pyghdet.

### 2.2 Empirical Data

The proposed method has been applied to two empirical datasets (the rattlesnake and the yeast datasets) [39, 40]. The rattlesnake dataset contains six subspecies of rattlesnake and two outgroup species and is known not to contain hybrid species. The yeast data set contains seven species of yeast and one outgroup and is known to contain one or more hybrid species [42]. The results are described in the following sections.

## 3 Results

### 3.1 Simulation Study

To evaluate the performance of the proposed statistic using the Cauchy Combination test, we plot the power curves for each of the scenarios (see the Appendix Section 6 for figures under all of the simulation settings). As a representative example, panel a) of Figure 3 shows power plots of the proposed test for the twenty-taxon tree with one ancient hybridization event (Figure 2e) with an *α* = 0.05. When *γ* = 0, these plots show that the power of the test is less than *α* = 0.05 across all the scenarios; i.e., under the null hypothesis of no hybridization, the power of the test is less than the selected level of significance. This result shows that the proposed test has the correct type-I error rate. Also, it is noticeable from the results that regardless of other parameters, as the length of the sequence increased, the power of the test also increased (setups with the length of sequence equal to 200k are the most powerful, and the power gradually decreases as the length decreases to 50k).

**Fig. 3:**
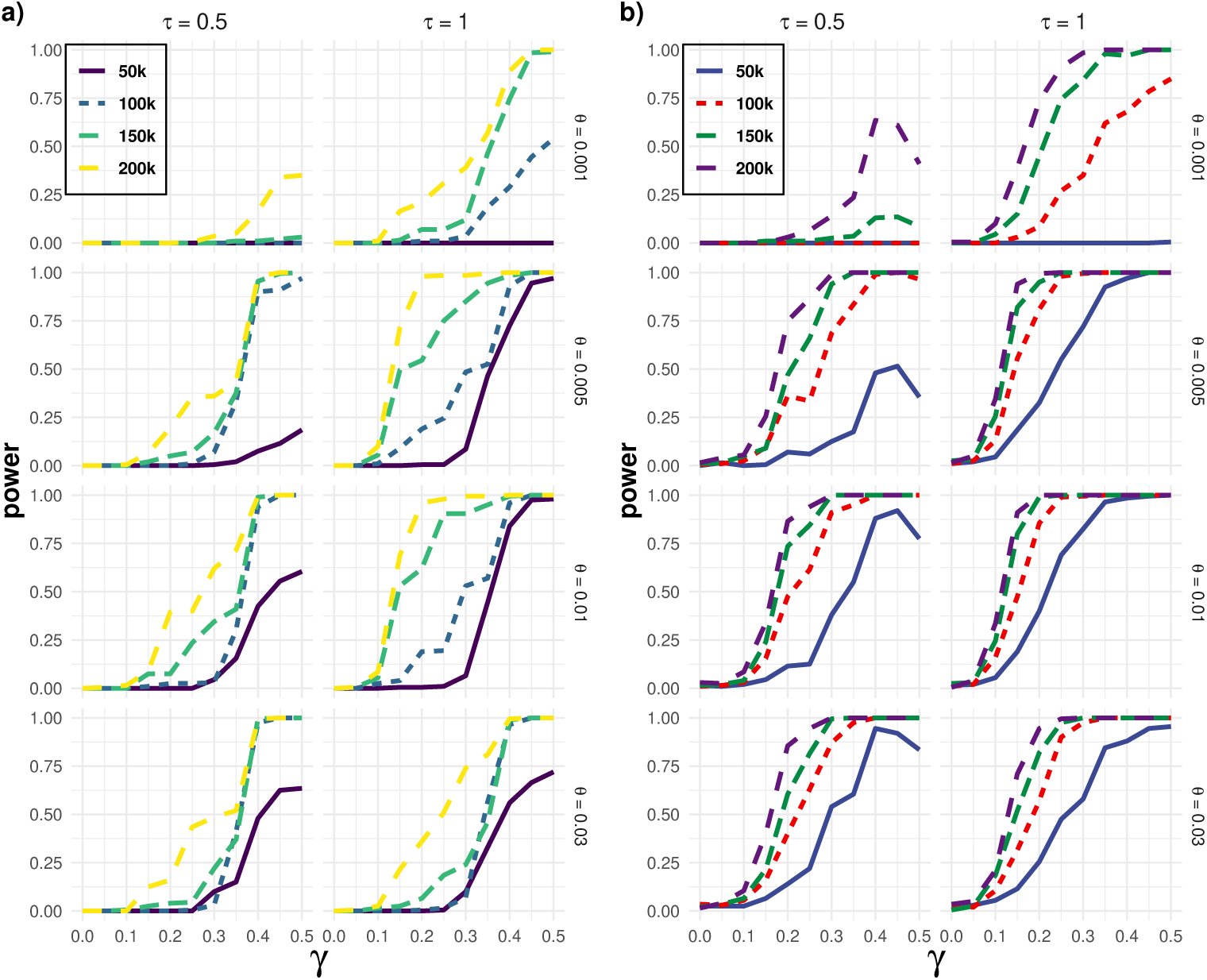
Results of the power simulations for the twenty-taxon hybrid species network with one ancient hybridization event in Figure 2e) using the Cauchy combination test (panel a) and the MCM test (panel b). Four different DNA sequence lengths (50*k*, 100*k*, 150*k*, and, 200*k*), four different effective population sizes (*θ* = 0.001, 0.005, 0.01, and, 0.03), two different branch lengths (*τ* = 0.50 and 1.0) has been considered. Here, *γ* represents the amount of genetic material contributed by each parent to the hybrid taxon.

The effect of branch length is also as expected. Holding everything else constant, tests for data with branch lengths equal to 1.0 are more powerful than those with 0.50. This result is expected because the prevalence of ILS increases in gene trees as branch length decreases, making hybridization harder to detect. We see a similar effect of the parameter *θ*, which controls the mutation rate (higher values correspond to more mutations). As expected, decreased value of *θ* results in decreased power to detect hybridization when other parameters are held constant. In addition, the power of the test is also increasing as the value of *γ* increases, regardless of other parameters. This result is also expected since a bigger value of *γ* is associated with a greater contribution from each parent, making it easier to detect hybridization. Overall, the test performs well. We can see that in almost all of the cases, the test reaches a high power (0.9 to 1.0) given a decent-sized sequence length (100k or more) when the value of *γ* is 0.25 or greater.

It is also worth noting that the test adheres to all the theoretical expectations. The only case when the proposed test does not perform well is when both the branch length and parameter *θ* are extremely small. As can be seen in the first panel of Figure 3, the test has extremely low power to detect hybridization when the branch lengths are 0.50 and *θ* = 0.001 even when the sequence length is high, and the value of *γ* is high. However, this is expected because, with branch lengths of 0.50, one should expect a large amount of ILS and an extremely low mutation rate as *θ* is extremely small. The combination of these two extreme situations makes it extremely hard to detect hybridization.

The performance of the global test using the MCM statistic proposed by Chen (2022) is similar to that of the test using the Cauchy combination test statistic proposed by Liu and Xie (2020). As a representative example, panel b) of Figure 3 shows power plots of the proposed test for the twenty-taxon tree with one ancient hybridization event (Figure 2e) with an *α* = 0.05. Figure 3 shows that the global test satisfies the theoretical expectations. We observed that the power of the test increases as the branch length (*τ*), length of DNA sequences (*N*), hybridization parameter (*γ*), and effective population size (*θ*) increases when the effect of each of these parameters are observed fixing all the other parameters.

We noticed that the power of the global test using the MCM test is slightly higher than the Cauchy combination test. For instance, the power of the test reaches 1.0 for a slightly lower value of *γ* when compared with the Cauchy combination test. Also, the power of the test using the MCM statistic is slightly higher in some cases when the length of the DNA sequence is small (e.g., *N* = 50*k*). However, the type-I error rate under the MCM test framework was greater than the chosen level of significance (*α*) in a few cases (7 out of 128 cases) (Figure 4), while the global test using the Cauchy combination framework maintained the true type-I error rate in all the scenarios.

**Fig. 4:**
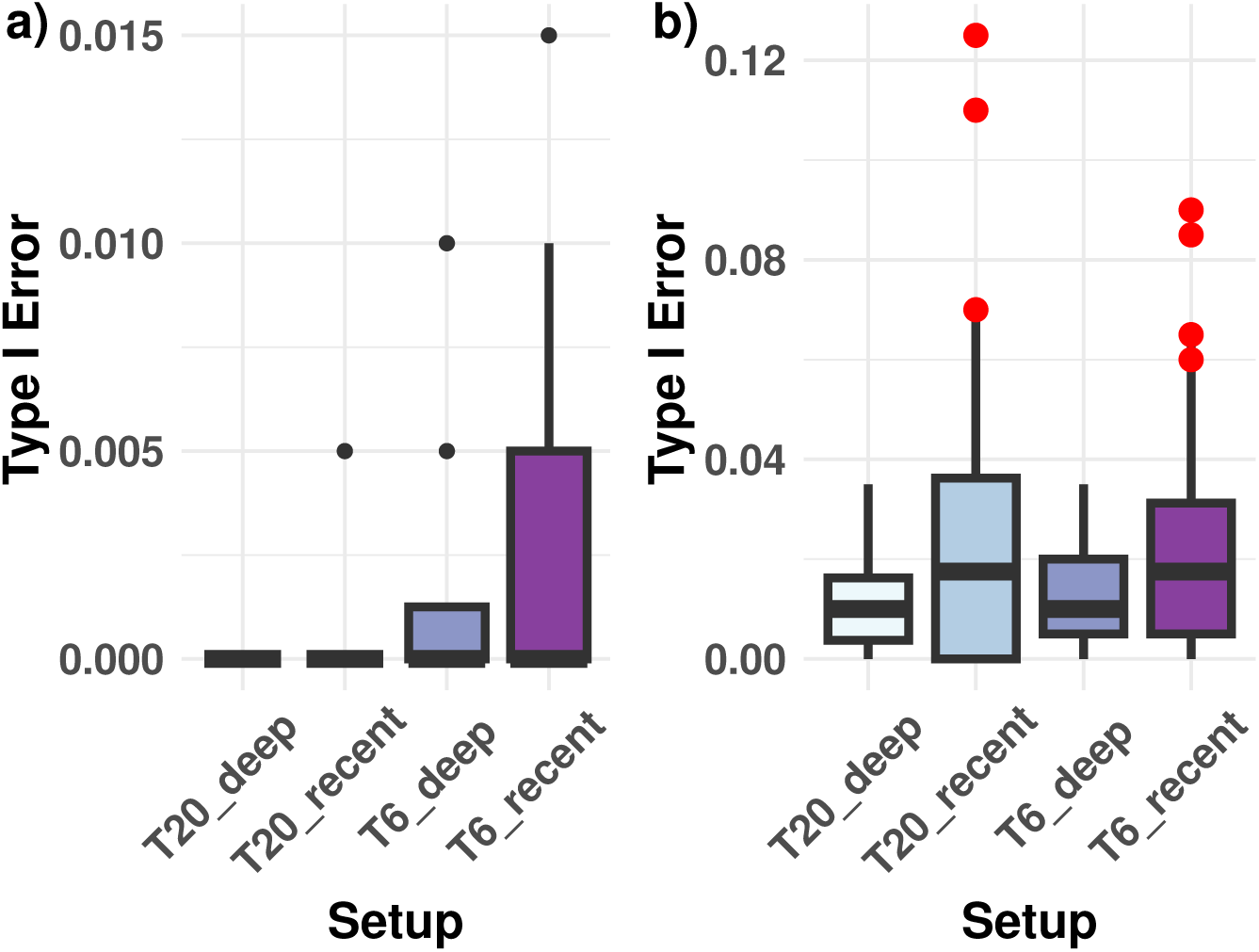
Type-I error rate under the Cauchy combination test (panel a) and the MCM test (panel b) for all the scenarios (T20_deep: twenty species with one ancient hybridization, T20_recent: twenty species with one recent hybridization, T6_deep: six species with one ancient hybridization, and T6_recent: six species with one recent hybridization). The red dots are situations where the type-I error rates were greater than the chosen level of significance (*α* = 0.05). For each setup, there are 32 simulated type-I error rates considering lengths of the DNA sequence alignment (50*k*, 100*k*, 150*k*, and, 200*k*), branch lengths (*τ* = 0.50 and *τ* = 1.0), and effective population sizes (*θ* = 0.001, 0.005, 0.01, and, 0.03). Each of these box plots consists of these 32 type-I error rates.

### 3.2 Empirical Data: Sistrurus Rattlesnakes

The dataset consists of six subspecies from two distinct rattlesnake species and two outgroup species, leading to a phylogeny with eight tips. Eighteen nuclear loci and one mitochondrial locus have been sequenced for a total of *N* = 8466 sites in the DNA sequence alignment. Our interest is to test the following null hypothesis:

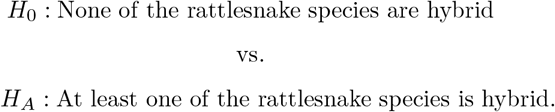

To perform this test at *α* = 0.05, we first created all possible subsets 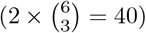 of the data with three ingroup species and one outgroup species. Once we have all the subsets, using the *H* statistic in Equation 4, we performed three hypothesis tests by considering one species as the hybrid and the other two as parental species using each of these subsets, totaling 3 × 40 = 120 individual tests. Finally, we used the resulting p-values from these individual tests in Equation 8 to compute the p-value of the global null hypothesis using the Cauchy combination test. We also calculated the global p-value for the MCM test using the p-values from the individual tests and the p-value from the Cauchy combination test in Equation 3. The p-values of both the Cauchy combination and MCM test were substantially larger than the chosen level of significance (*p_c_* > 0.99 and *p_mcm_* > 0.99). Hence, we failed to reject the null hypothesis and concluded that there were no hybrid species within the six subspecies of rattlesnake. This finding is consistent with other analyses using this dataset [39, 40].

### 3.3 Empirical Data: Yeast

The yeast data set of Rokas et al. (2003) contains seven Saccharomyces species, *S. cerevisiae* (Scer), *S. paradoxus* (Spar), *S. mikatae* (Smik), *S. kudriavzevii* (Skud), *S. bayanus* (Sbay), *S. castellii* (Scas), *S. kluyveri* (Sklu), and the outgroup fungus *Candida albicans* (Calb). The sequences come from 106 genes, totaling 127, 026 sites in the DNA sequence alignment. Using the dataset, we have 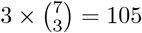 quartet trees, each with two parents and one potential hybrid from the seven species and the outgroup. We aim to test the following null hypothesis:

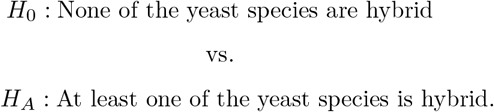

To perform this test, we calculated the *H* statistic (Equation 4) for all of the 105 possible quartet trees, along with p-values for all the individual tests. Finally, using the p-values from all the individual tests, we computed the p-value (*p_c_*) of the Cauchy combination test for the global null hypothesis using Equation 8. We also computed the global p-value (*p_mcm_*) of the MCM test using p-values from all the individual tests and the p-value from the Cauchy combination test in Equation 3. The p-value for the global test was found to be less than 0.001 for both the Cauchy combination and the MCM test. Hence, we have enough evidence to reject the null hypothesis and conclude that the yeast dataset contains at least one hybrid species. A network analysis of the yeast dataset by Yu et al. (2011) found several hybrid species among the seven species. Hence, our result is consistent with Yu et al. (2011).

## 4 Discussion

We have proposed two methods for assessing whether a data set contains one or more hybrid species that take incomplete lineage sorting into account. Based on DNA sequence data on a set of taxa, we propose our test as the first step before inferring a phylogenetic tree or network. One of the key benefits of the proposed method is its speed. The computation time of the proposed test is almost negligible since we used observed site pattern frequencies to calculate the test statistic. The speed makes the test feasible for genome-scale data sets with a large number of taxa. On one hand, individual tests are based on observed site patterns derived from a multinomial distribution, which allows a normal distribution to be derived from the asymptotic distribution of the site patterns for the test statistic. On the other hand, the null distribution of the global test was standard Cauchy, leading to a straightforward method for calculating the p-value.

Our simulation studies show that the methods are powerful to detect any presence of hybridization events even when the hybridization event is close to the root. Results show that deep or ancient hybridization events are harder to detect compared to recent hybridization events. However, if the branch length and length of the DNA sequence are large enough, the proposed method can still detect the presence of hybridization with high power. The proposed methods are extremely efficient: for six taxon networks with 250, 000 sites, the whole process can be completed within 25 seconds, while for a data set with 20 taxa and 1 million sites, the whole process can be completed within 10 minutes. This shows that the proposed methods are extremely scalable for large data sets with a large number of sites. It is also possible to use parallel computing for the individual tests and then combine the results at the end if more time efficiency is needed.

The methods are based on the Hils statistic proposed by Kubatko and Chifman (2019) in combination with two combination tests (the Cauchy combination test, and the MCM test). We notice that the type I error rate under the MCM test is sometimes slightly larger than the chosen level of significance, and the Cauchy combination test is more conservative. Thus, there is a possibility of improvement by making the type I error rate smaller for the MCM test while maintaining the same power. There are also opportunities to try to reduce the number of individual tests needed to perform the global test. As we are only interested to know whether one or more hybrid taxa are present in a data set, it is reasonable to perform one test per quartet tree to determine if there are any hybrid species in that set and finally combine the results.

## 5 Conclusion

In many areas of biological research, it is central to understanding the relationships between species. There are two primary ways to represent such relationships: a species-level phylogeny and a species network. Therefore, depending on the data, it is necessary to decide which analysis method to use. In doing so, we face several challenges, such as computational efficiency, modeling the data at the gene level as well as species level, and developing methods to account for potential evolutionary mechanisms (e.g., incomplete lineage sorting, gene duplication, hybridization) that generate incongruence between individual gene histories. The multispecies coalescence model is commonly used to model incomplete lineage sorting and can be used to generate gene trees within a species tree. We considered coalescence theory while generating data to detect possible hybridization events in a given data set. We demonstrated that the proposed method is powerful in detecting hybridization events in several possible situations via simulation. We also demonstrated the method is capable of detecting hybridization events in empirical data. It is also notable that our method is extremely efficient and thus can handle genome-scale data with a large number of taxa. Thus, the proposed method provides researchers with an efficient way of detecting hybridization events in a data set and formally deciding whether a tree or a network analysis is more appropriate for a given data set.

## 6 Appendix

**Fig. 5:**
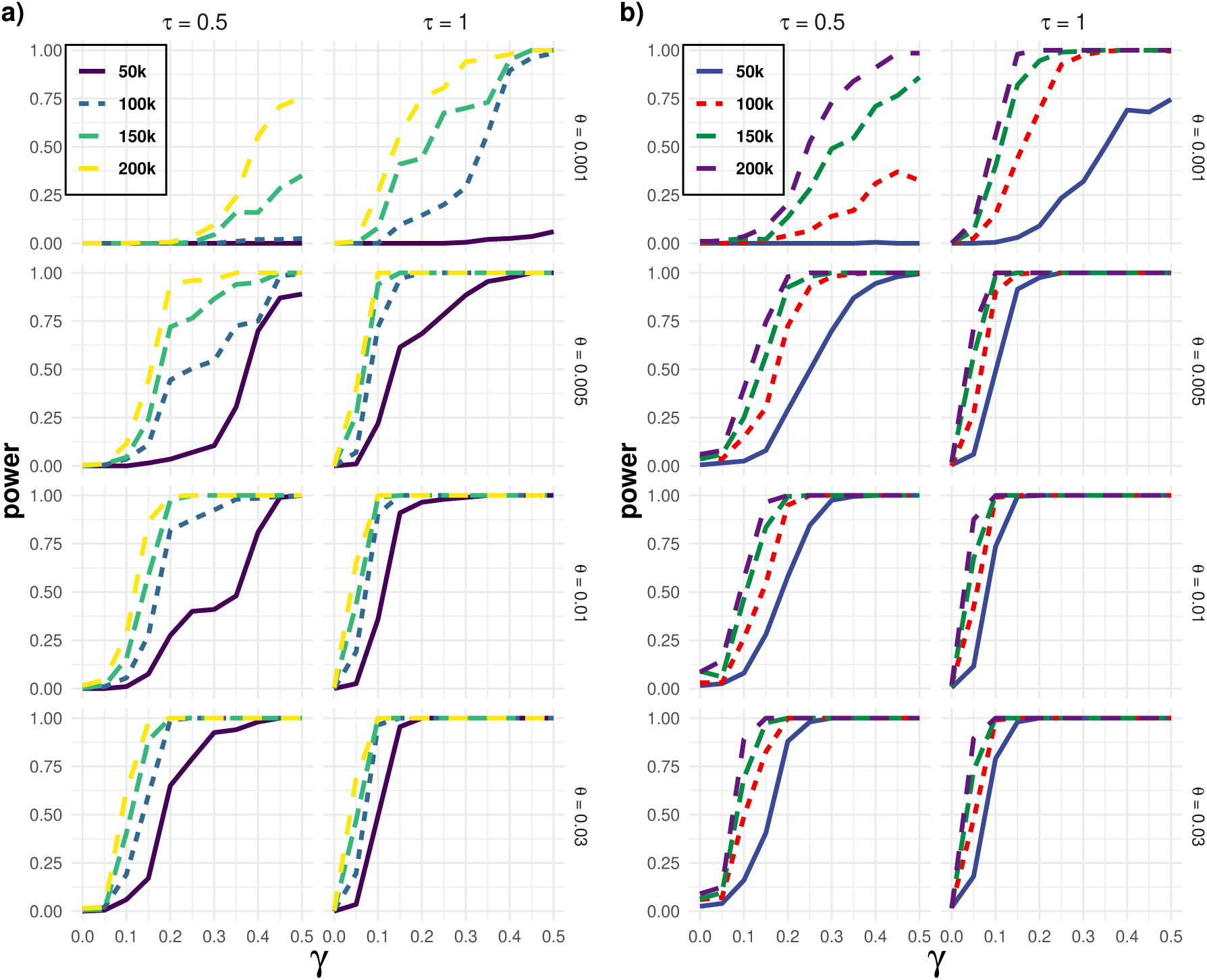
Results of the power simulations for the six-taxon hybrid species network with one recent hybridization event in Figure 2a) using the Cauchy combination test (panel a) and the MCM test (panel b). Four different DNA sequence lengths (50*k*, 100*k*, 150*k*, and, 200*k*), four different effective population sizes (*θ* = 0.001, 0.005, 0.01, and, 0.03), two different branch lengths (*τ* = 0.50 and 1.0) has been considered. Here, *γ* represents the amount of genetic material contributed by each parent to the hybrid taxon.

**Fig. 6:**
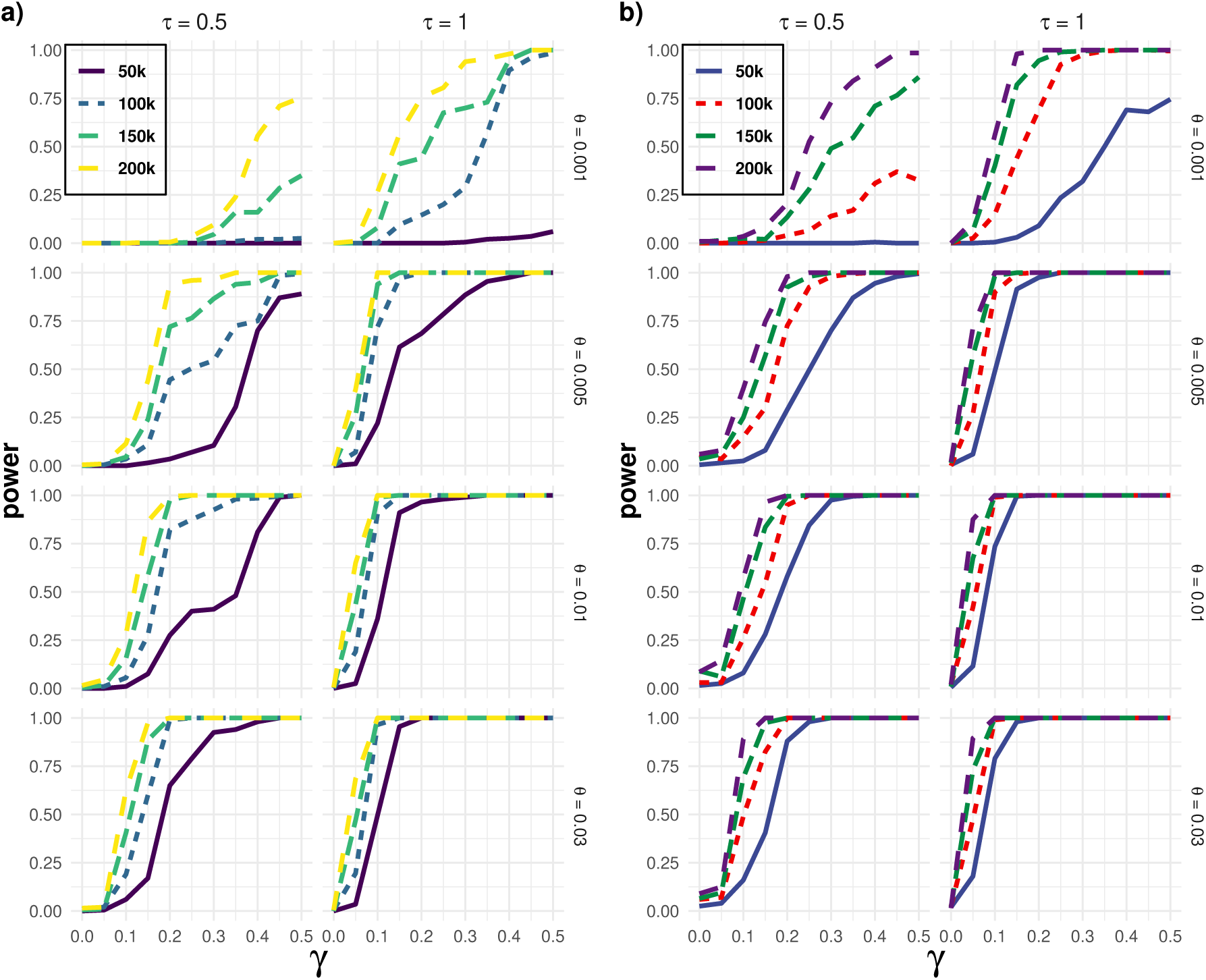
Results of the power simulations for the six-taxon hybrid species network with one ancient hybridization event in Figure 2b) using the Cauchy combination test (panel a) and the MCM test (panel b). Four different DNA sequence lengths (50*k*, 100*k*, 150*k*, and, 200*k*), four different effective population sizes (*θ* = 0.001, 0.005, 0.01, and, 0.03), two different branch lengths (*τ* = 0.50 and 1.0) has been considered. Here, *γ* represents the amount of genetic material contributed by each parent to the hybrid taxon.

**Fig. 7:**
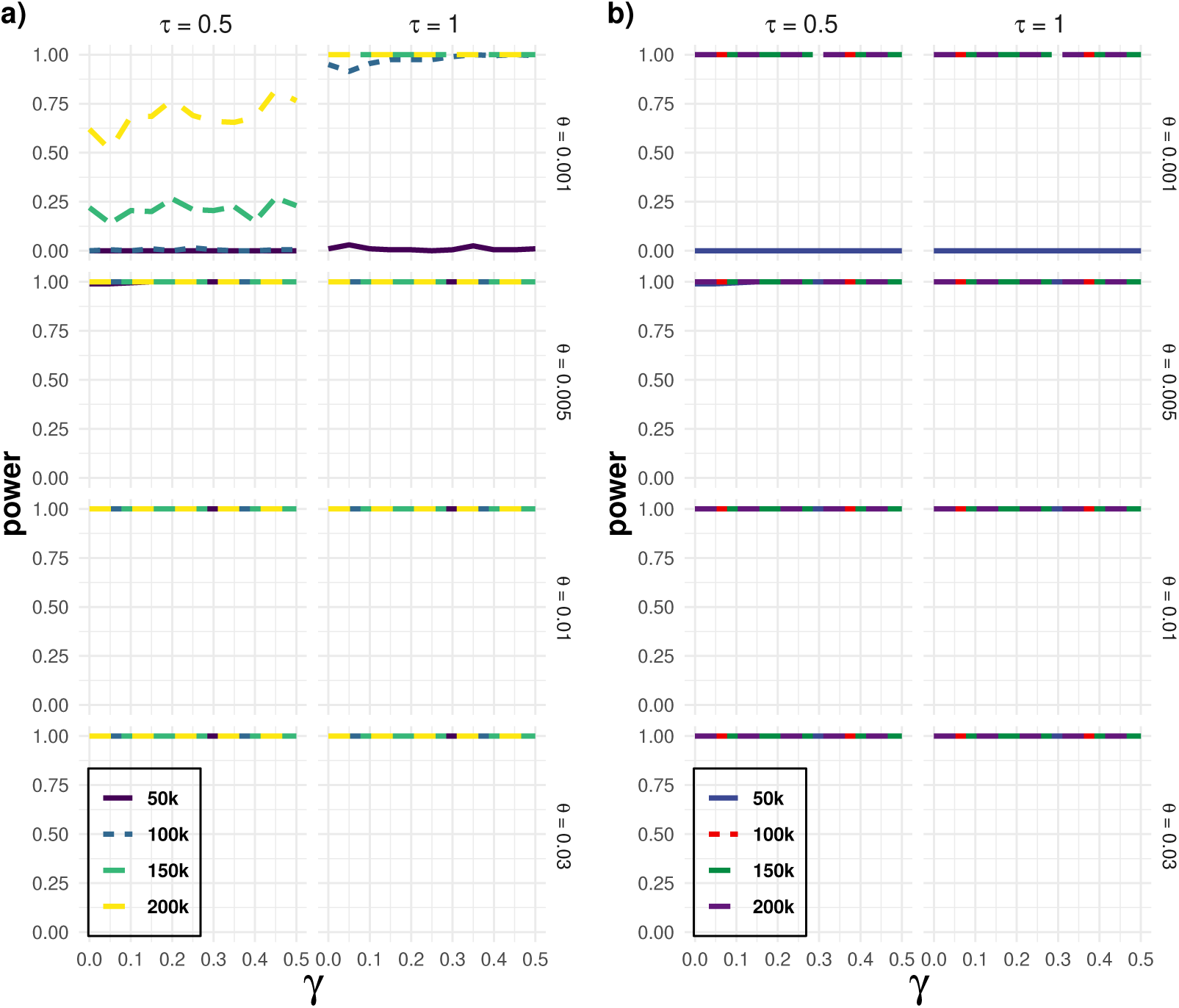
Results of the power simulations for the seven-taxon hybrid species network with one ancient and one recent hybridization events in Figure 2c) under the Cauchy combination test framework (panel a) and MCM test framework (panel b). Four different DNA sequence lengths (50*k*, 100*k*, 150*k*, and, 200*k*), four different effective population sizes (*θ* = 0.001, 0.005, 0.01, and, 0.03), two different branch lengths (*τ* = 0.50 and 1.0) has been considered. Here, *γ* represents the amount of genetic material contributed by each parent to the hybrid taxon *h*_1_ while fixing *γ*_2_ at 0.50 for hybrid taxon *h*_2_.

**Fig. 8:**
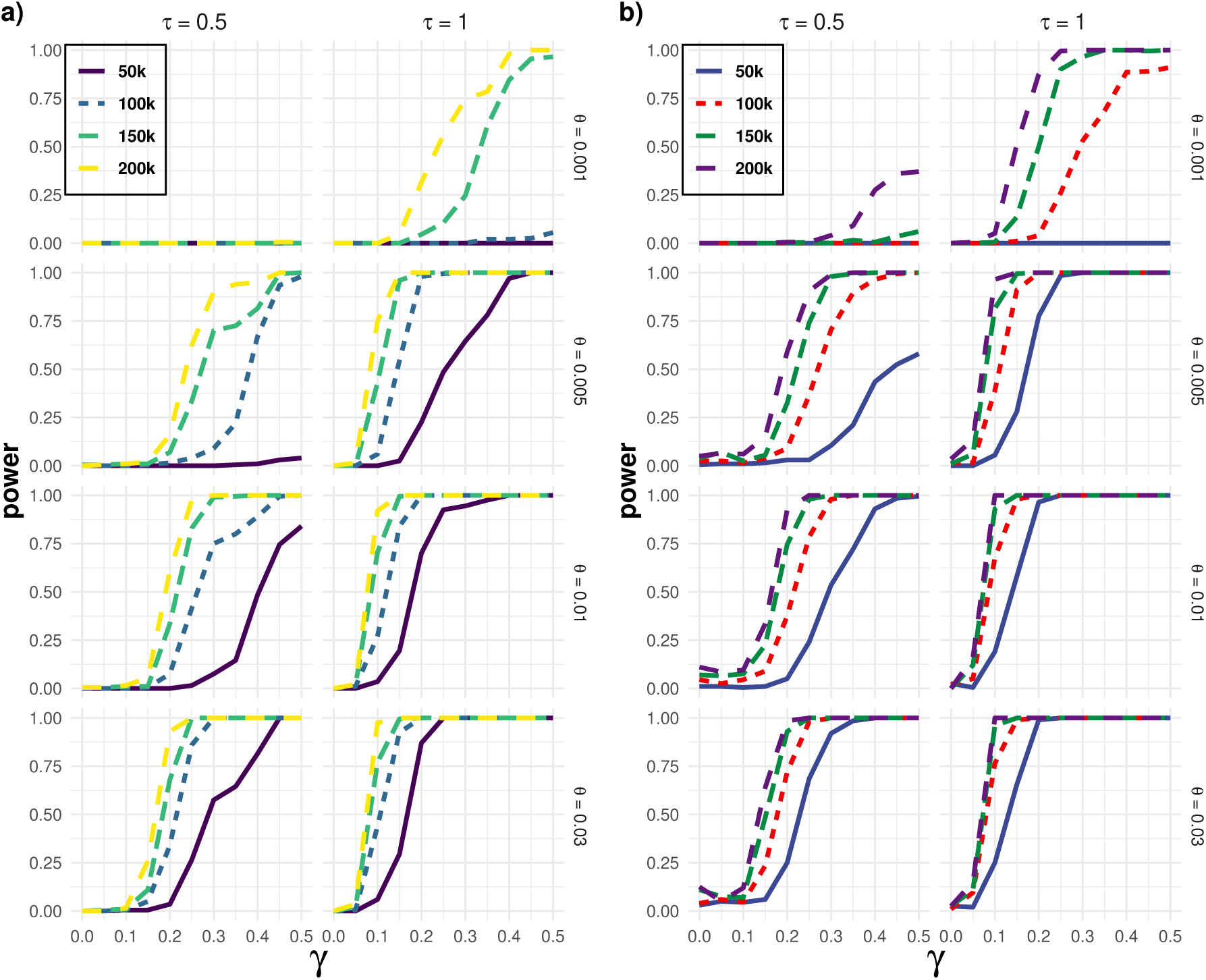
Results of the power simulations for the twenty-taxon hybrid species network with one current hybridization event in Figure 2d) using the Cauchy combination test (panel a) and the MCM test (panel b). Four different DNA sequence lengths (50*k*, 100*k*, 150*k*, and, 200*k*), four different effective population sizes (*θ* = 0.001, 0.005, 0.01, and, 0.03), two different branch lengths (*τ* = 0.50 and 1.0) has been considered. Here, *γ* represents the amount of genetic material contributed by each parent to the hybrid taxon.

